# Measurement force, speed and post-mortem time affect the ratio of CNS grey to white matter elasticity

**DOI:** 10.1101/2025.01.30.635764

**Authors:** Julia M. Becker, Alexander K. Winkel, Eva Kreysing, Kristian Franze

## Abstract

For several decades, many attempts have been made to characterise the mechanical properties of grey and white matter, which constitute the two main compartments of the central nervous system (CNS), with various methods and contradictory results. In particular, the ratio of grey-to-white-matter elasticity is sometimes larger than 1 and sometimes smaller; the reason for this apparent discrepancy is currently unknown. Here, we exploited atomic force microscopy (AFM)-based indentation measurements to systematically investigate how the measurement force, measurement speed, post-mortem interval and temperature affect the measured elasticity of spinal cord tissue, and in particular the ratio of grey-to-white-matter elasticity (*K_g_/K_w_*). Within the explored parameter space, increasing measurement force and speed increased the measured elasticity of both grey and white matter. However, *K_g_/K_w_* declined from values as high as ∼5 at low forces and speeds to ∼1 for high forces and speeds. *K_g_/K_w_* also strongly depended on the anatomical plane in which the measurements were conducted and was considerably higher in transverse sections compared to longitudinal sections. Furthermore, the post-mortem interval impacted both the absolute measured tissue elasticity and *K_g_/K_w_*. Grey matter elasticity started decreasing ∼3 hours post-mortem until reaching a plateau after ∼6 hours. In contrast, white matter elasticity started declining from the beginning of the measurements until ∼6 hours post-mortem, when it also levelled off. As a result, *K_g_/K_w_* increased until ∼6 hours post-mortem before stabilising. Between 20°C and 38°C, both grey and white matter elasticity decreased at a similar rate, without affecting *K_g_/K_w_*. We have thus identified differences in the response of grey and white matter to varying strains and strain rates, and the post-mortem interval, and excluded temperature as a factor affecting *K_g_/K_w_*. These differential responses likely contribute to the contradictory results obtained with different methods working in different strain regimes.

**Statement of significance:** We here showed that the mechanical response of CNS grey and white matter to an applied force differentially depends on measurement parameters such as the speed and magnitude of the applied forces, the post-mortem interval, as well as the anatomical axis along which measurements are conducted. These results broaden our understanding of CNS mechanics and pave the way for better and more targeted experimental design of future experiments. Ultimately, they may help to reconcile seemingly contradictory results in the literature concerning the ratio of grey-to-white-matter elasticity.

## Introduction

The mechanical properties of central nervous system (CNS) tissue are critical for regulating diverse neuronal and glial cell functions (1). Tissue stiffness, for example, contributes to developmental processes, such as axon path finding (2, 3) and brain folding (4), to tissue maintenance during adult neurogenesis (5), as well as to pathological processes, such as brain tumour chemo-resistance (6) or axonal remyelination (7). Knowledge about mechanical CNS tissue properties, which are highly dynamic and change during development, ageing, and pathological processes, is therefore important for our understanding of normal and dysregulated CNS cell function.

For over half a century, various methods have been exploited to measure CNS mechanics in health and disease, resulting in differing and sometimes even contradictory findings. In particular, there is still no consensus regarding mechanical differences between grey matter, which mostly contains neuronal cell bodies and synapses, and white matter, which is predominantly built up by myelinated axon tracts. The ratio of grey-to-white-matter tissue elasticity (*K_g_/K_w_*) is not only important for cortical folding (4) but also in the context of mechanical trauma, as the relative mechanical properties of grey and white matter affect the stress and strain patterns resulting from traumatic deformations (8). Yet, in some reports, grey matter is stiffer than white matter (9–13), while in others, white matter is stiffer than grey matter (14–17), and there are publications where both tissue types are mechanically similar (18, 19). These findings somewhat cluster according to the methodologies used to measure tissue mechanics. Studies employing atomic force microscopy (AFM) (10–13), which applied forces in the ∼nN regime at a frequency on the order of ∼1 Hz, usually find *K_g_/K_w_* > 1, whereas studies using nanoindentation testing (14–16) or magnetic resonance elastography (MRE) (17), which apply comparatively larger forces (∼hundreds of µN) or forces at higher frequencies (∼50 – 100 Hz), respectively, commonly find *K_g_/K_w_* < 1.

Although it is known that many parameters influence overall CNS tissue elasticity, such as the applied strain amplitude (i.e. the amount of deformation) (20, 21), the strain rate (i.e. the speed of deformation) (10, 15), the ‘freshness’ of the sample (i.e., the post-mortem time) (22, 23), and the temperature (24, 25), their effect on the grey-to-white-matter elasticity ratio *K_g_/K_w_* has not yet been systematically investigated.

Here, we used atomic force microscopy (AFM)-based indentation tests to determine the reduced apparent elastic modulus 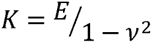 (a measure of elasticity, with *E* being the Young’s modulus and *v* the Poisson’s ratio) of fresh rat spinal cord sections, which are characterised by clearly defined grey and white matter areas. We conducted representative elasticity measurements across entire sections, systematically varying the measurement force (i.e. strain), measurement speed (i.e. strain rate), anatomical plane (i.e. directionality of the force application relative to the orientation of the tissue), post-mortem interval, and measurement temperature. We found that, while temperature did not affect the grey-to-white-matter elasticity ratio *K_g_/K_w_*, force, speed, anatomical plane and the post-mortem interval did.

## Materials and methods

For ease of reading, general protocols are presented first, followed by details and deviations for individual experiments. All reagents were purchased from Sigma-Aldrich, unless stated otherwise.

### Artificial cerebrospinal fluid (aCSF) solutions

Artificial cerebrospinal fluid (aCSF) solutions established for electrophysiology (26) were used to ensure optimal tissue quality and prepared fresh every day. Slicing aCSF contained 5 mM ethyl pyruvate, 26 mM choline bicarbonate, 2 mM NaOH (Fisher Scientific), 2 mM kynurenic acid, 191 mM sucrose, 20 mM glucose, 1 mM (+)-sodium L-ascorbate, 3 mM myo-inositol, 0.75 mM potassium gluconate, 1.25 mM KH_2_PO_4_, 4 mM MgSO_4_ and 1 mM CaCl_2_ (Fluka) in autoclaved ddH_2_O. Measuring aCSF contained 5 mM ethyl pyruvate, 15 mM glucose, 1 mM (+)-sodium L-ascorbate, 3 mM myo-inositol, 121 mM NaCl, 3 mM KCl, 1.25 mM NaH_2_PO_4_, 25 mM NaHCO_3_, 1.1 mM MgCl_2_ and 2.2 mM CaCl_2_ (Fluka) in autoclaved ddH_2_O. Both aCSF solutions were bubbled with carbogen (95% O_2_/5% CO_2_) for at least 30 mins before and throughout the experiments. Slicing aCSF was kept on ice before and during use, measuring aCSF was kept at room temperature. The resulting pH was 7.20 – 7.33 for the slicing aCSF and 7.34 for the measuring aCSF.

### Animals and sample preparation

Details about the rats used in this study are shown in Table 1. To reduce animal numbers, spare rats from other research groups were used when available; otherwise, rats were purchased from Charles River and Envigo. All work involving animals was conducted in keeping with the Animals (Scientific Procedures) Act 1986 and approved by the relevant authorities according to the University of Cambridge Animal Welfare and Ethical Review Body (AWERB) processes.

**Table 1:** Animals used for experiments.

| Experiment | Strain | Sex & numbers | Median weight<br>(range) [g] | Estimated age<br>[weeks] | Sectioning plane &<br>vertebral level |
| --- | --- | --- | --- | --- | --- |
| Force and speed | Lister Hooded | Female (N = 13);<br>Male (N = 9) | 219 (165 - 283) | 6 – 10 | Transverse (C5);<br>Sagittal (C4 – C6);<br>Horizontal (C4 – C6) |
| Post-mortem interval | Wistar; Wister<br>Han | Female (N = 11) | 330 (282 - 509) | 46 – 49 | Transverse (C5) |
| Temperature | Lister Hooded | Female (N = 6) | 215 (200 - 226) | 9 – 10 | Transverse (C5) |

Rats were euthanised with an overdose of pentobarbital (2000 mg/kg i.p.) under isoflurane anaesthesia. Typical times required for individual steps of the dissection procedure relative to the time of death are indicated in brackets. A thoracotomy was performed and a shortened intravenous catheter was placed through the left atrium into the proximal aorta and stitched in place (∼3 mins), and immediate perfusion with ice-cold slicing aCSF at 15 – 16 ml/min was started. A dorsal midline incision was made and the cervical spinal cord of the vertebral segments C4 to C6 was exposed via a laminectomy and gently removed from the body (∼18 mins). Dissection was continued in a perfused dish (5 ml/min) on ice and the dura mater and leptomeninx were carefully removed (∼35 mins). Samples were embedded in 4% low gelling temperature agarose in PBS and sectioned on a Leica VT1000S vibratome in slicing aCSF on ice with a Gillette Platinum blade. Section thickness was set to 999 µm, forward speed to 50 µm/s, vibration frequency to 60 Hz and vibration amplitude to 1 mm. Sections were immediately transferred to bubbled measuring aCSF at room temperature (∼50 to 75 mins). When sectioning the horizontal or sagittal planes, care was taken to select tissue sections for AFM measurements where the location of the grey-white matter boundary did not change much through the thickness of the section. The section chosen for AFM was transferred to a tissue culture dish (Ø 40 mm, TPP), held in place by small dots of cyanoacrylate glue between the dish and the agarose surrounding the sample, and submerged in measuring aCSF.

### AFM setup and measurements

Force-distance curves were acquired with our custom-built AFM setup in spectroscopy mode. The setup consists of a CellHesion® 200 AFM head on a motorized precision stage with a Vortis Advanced Control Station (all JPK) and an AxioZoom.V16 upright microscope (Zeiss) with an Andor camera (Zyla sCMOS), mounted on an air table (Newport, RS 2000^TM^). The dish with the tissue sample was placed in a PetriDishHeater (JPK) set to 34°C, which resulted in a temperature of ∼32.5°C near the sample. To assess the effect of temperature, the samples were instead placed inside a BioCell (JPK), using setpoint temperatures of 20°C, 24.5°C, 29°C, 33.5°C and 38°C in either ascending or descending order (N = 3 animals each). Samples were given sufficient time (∼15 mins) to equilibrate at each new temperature level before measurements were started. In all cases, bubbled measuring aCSF was continuously exchanged with a perfusion pump throughout the experiment at a rate of 0.72 ml/min.

For the majority of experiments, we used Arrow^TM^ TL1 tipless silicon cantilevers (NanoWorld) with a spring constant of 0.083 – 0.093 N/m. For experiments investigating the effect of force and speed, an Arrow^TM^ TL1 cantilever with a spring constant of 0.374 N/m (N = 2 animals) and TL-FM cantilevers (Nanosensors) with spring constants of 1.849 – 2.757 N/m (N = 20 animals) were used instead to allow for higher indentation forces. Cantilever spring constant was determined contact-free in air with the in-built thermal noise method of the JPK SPMControl software. Spherical polystyrene beads of 44.65 μm radius (microParticles GmbH) were glued to the cantilever tip using M-Bond 610 (Agar Scientific). Cantilever sensitivity was determined on a glass slide in buffer before each experiment. AFM measurements were conducted at a constant indentation speed of 20 μm/s in closed loop with a force of 30 nN at a sample rate of 1000 Hz. Experiments investigating the effect of force and speed were conducted with varying combinations of indentation forces of 30, 90, 150, 300, 600, 1200 and 1500 nN and indentation speeds of 20, 100, 400, 800 and 1200 µm/s in closed loop, and sample rates were chosen to ensure a minimum of 2500 points on the extend part of the recorded force-distance curve. AFM measurement grids containing many individual measurement points, also called “maps”, were tailored to each sample’s geometry using a custom-written MATLAB (The MathWorks) routine in order to cover the sample in a representative way. Generally, maps were measured in mediolateral direction; to investigate the effect of force and speed, various measurement directions were used. Table 2 contains the start and end time of AFM measurements for individual experiments and the grid resolutions used. To examine the effect of post-mortem time and temperature, the same map was repeatedly remeasured, either after a certain amount of time had elapsed or after sample temperature had been changed, respectively. In the former case, maps were repeatedly measured every ∼30 or ∼60 minutes. Maps which covered both halves of the spinal cord took ∼90 mins to complete.

**Table 2:** Start and end time and resolution of AFM measurements.

| Experiment | Measurement start<br>[hh:mm post-mortem] | Measurement end<br>[hh:mm post-mortem] | Grid resolution<br>[μm] |
| --- | --- | --- | --- |
| Force and speed | 1:30 – 2:30 | 3:05 – 8:35 | 50 – 200 |
| Post-mortem interval | 1:30 – 2:04 | 6:27 – 11:06 | 178 – 260 |
| Temperature | 2:10 – 2:48 | 5:43 – 6:11 | 182 – 201 |

### AFM data analysis

All AFM data analysis was carried out with MATLAB (The MathWorks). A custom-written script (10) was used to determine the reduced apparent elastic modulus *K* from each force-distance curve after determining the contact point with a fitting algorithm. All elasticity values are reported as *K* rather than the Young’s modulus *E* to avoid assumptions about the value of Poisson’s ratio *v* (*K* = *E*/(1-*v*^2^). To obtain *K*, the Hertz model 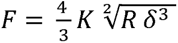 (27) was fitted to the curve data (*R* is bead radius; *F* is exerted force; 12 is indentation depth) using a custom-written script. The use of the Hertz model for a paraboloid indenter was considered appropriate for the analysis of the post-mortem time and temperature data acquired with a spherical indenter, as more than two thirds of all measurements satisfied 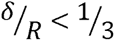, where both indenter geometries are similar. For data analysis concerning the effect of force and speed, the in-built Sneddon model for a spherical indenter (28, 29) from the JPK Data Processing Software was used instead to provide greater accuracy for higher indentation depths. Using an overview image of each sample with an overlay of the measurement grid, data points were manually segmented into white and grey matter. Custom-written software for AFM and AFM data analysis is available from https://github.com/FranzeLab/AFM-data-analysis-and-processing/tree/Batchforce_1.1/Batchforce and https://github.com/FranzeLab/AFM-data-analysis-and-processing/tree/master/JuliaBeckerThesis. All raw data force-indentation curves used for analysis are available online (DOI: 10.5281/zenodo.14630529).

To obtain the stiffness *k* for comparison with the apparent reduced elastic modulus *K*, a slope was fitted to the force-indentation data in the range of 90 – 100% exerted force using the JPK Data Processing Software.

### Statistical and regression analysis

Statistical analysis was carried out with Prism 9 or 10 for macOS (GraphPad Software). The significance level was α = 0.05. To assess whether AFM measurements themselves altered the tissue’s mechanical properties, a two-tailed one-sample t-test against a hypothetical value of 1 was used. To assess the effect of directionality, ANOVA was used to compare *K* values and the *K_g_/K_w_* ratio obtained in different anatomical planes with the same parameter combinations for force and speed, followed by Tukey’s multiple comparisons test for individual group comparisons. For this analysis, only parameter combinations were assessed for which data was available for all three planes and at least three animals per plane. To assess the effect of repeated AFM measurements, a two-tailed unpaired t-test was used on log-transformed data. For regression analysis concerning the effect of temperature, data points from all maps of all animals were pooled. Each data point constituted the median apparent elastic modulus over the median temperature of one elasticity map in one animal. For the *K_g_/K_w_* ratio, the ratio of the median grey to the median white matter apparent elastic modulus over the mean of the median grey and white matter temperature was used. An extra-sum-of-squares F test was used to compare a linear regression model to a horizontal line. To investigate the relationship between the apparent reduced elastic modulus K and the stiffness k, correlation analysis and linear regression was conducted with MATLAB (The MathWorks).

## Results

To minimize cell damage and maximise sample viability, care was taken to dissect the tissue as carefully and quickly as possible, while constantly supplying it with oxygenated buffer solutions optimised for rodent CNS tissue (26) at specific temperatures (for details, see Material and methods). AFM measurements started at approximately 1.5 – 2 hrs post-mortem. Throughout the study, we used AFM cantilevers with spherical beads of 44.65 μm radius glued to their tips to exert forces on spinal cord tissue sections.

### Force and speed of AFM measurements strongly affect the measured tissue elasticity and K_g_/K_w_

Atomic force microscopy permits spatially resolved mechanical probing of a tissue section. Force-distance curves obtained during an experiment can be fitted with the Sneddon or Hertz models (see methods) to obtain the measured tissue elasticity, i.e. the reduced apparent elastic modulus *K*. AFM also allows for systematic variation of key measurement parameters, i.e. force and speed, which affect the measured *K* values (Figure 1).

**Figure 1:**
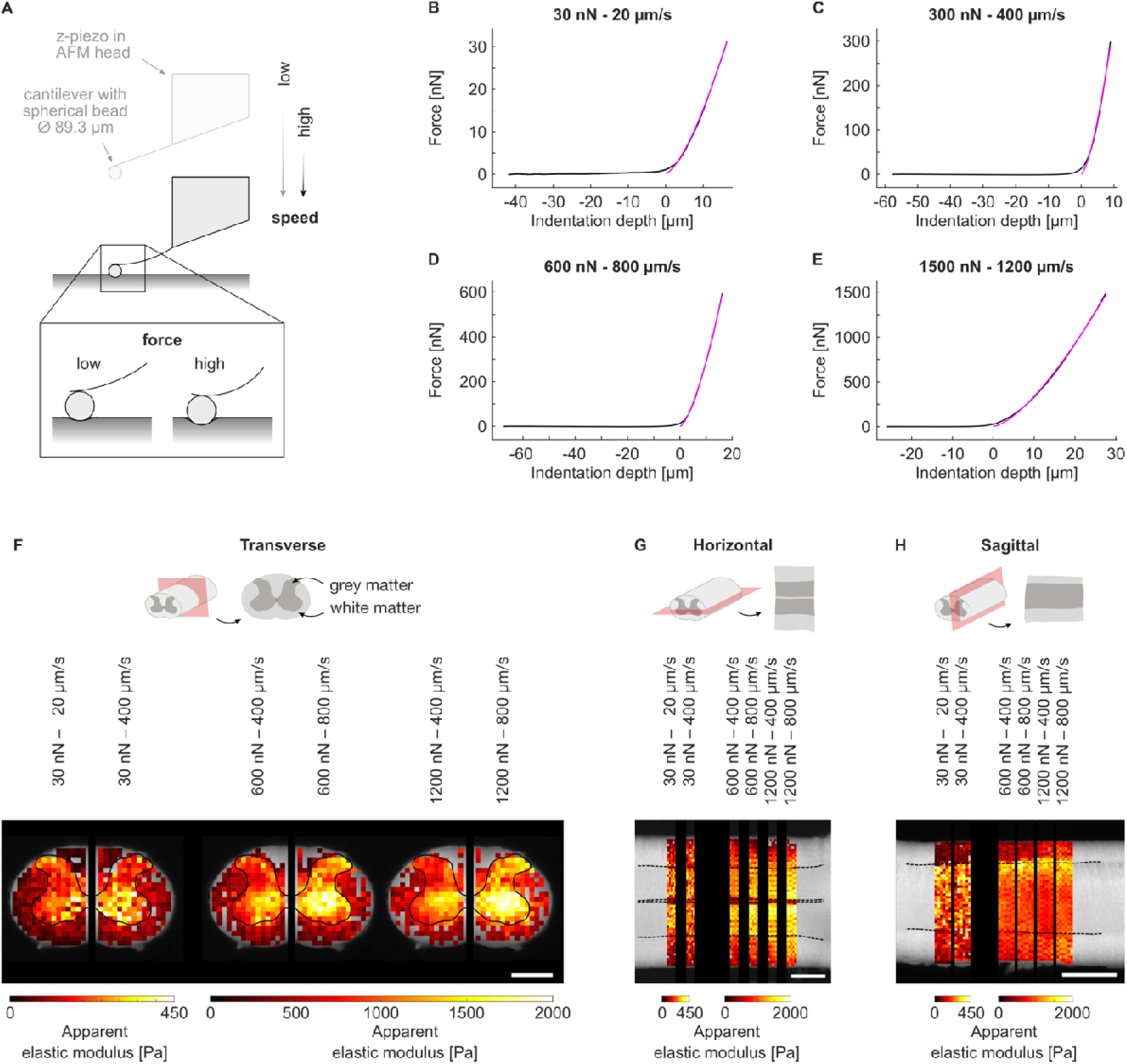
Choice of AFM measurement parameters force and speed affects the apparent reduced elastic modulus *K*. (**A**) Schematic of atomic force microscopy (AFM) setup. (**B-E**) Example force-distance curves (black) obtained from AFM measurements with different parameter combinations. The Sneddon model (pink) was fitted to the extend parts of the force-distance curves to obtain the respective apparent reduced elastic modulus *K*. (**F-H**) Schematics of the (**F**) transverse, (**G**) horizontal and (**H**) sagittal planes (grey matter inside: dark grey, white matter outside: light grey, anatomical plane: red). For each anatomical plane, six AFM elasticity heatmaps are shown which were acquired on the same example tissue section with different combinations of measurement forces and speeds. Different parameter combinations yield different apparent reduced elastic moduli on the same tissue section. Elasticity heatmaps are overlaid on a brightfield image of the measured tissue section. Images are broken for better visual segregation of the different parameter combinations. Each pixel represents one measurement. Dashed black lines: grey-white-matter boundary. Scale bars: 1000 µm.

To assess how force and speed affect *K*, which effectively is a measure of tissue stiffness, in different anatomical planes, we conducted AFM indentation measurements on transverse, horizontal and sagittal spinal cord tissue sections and systematically varied both measurement force *F* and measurement speed *s* within the limits dictated by the setup’s capabilities over two orders of magnitude (*F* = 30 – 1500 nN and *s* = 20 – 1200 µm/s) (Figure 2).

**Figure 2:**
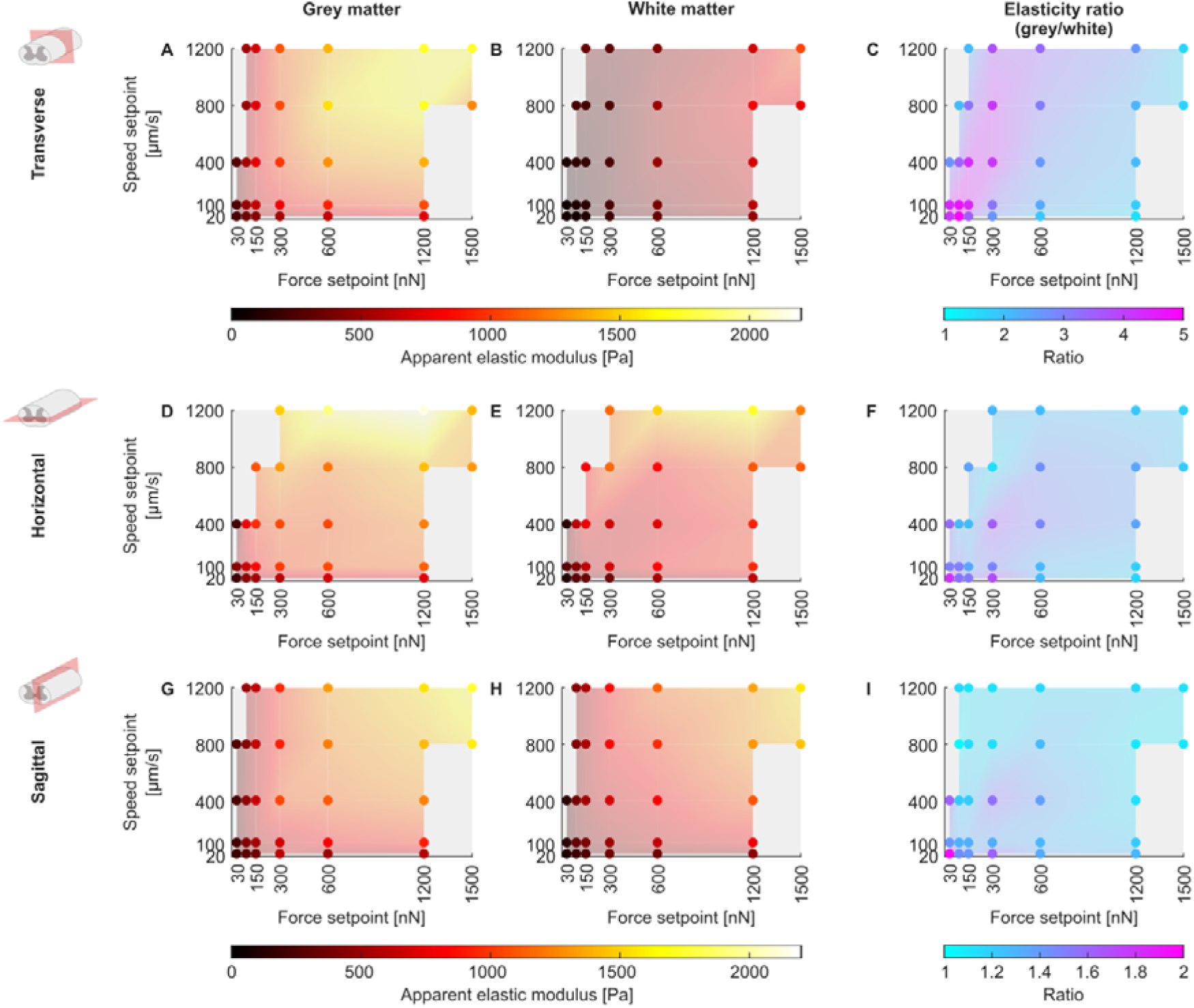
Increasing force and speed of AFM measurements increases the measured elasticity of grey and white matter but decreases the ratio of grey-to-white-matter elasticity. Apparent elastic moduli *K* for (**A, D, G**) grey and (**B, E, H**) white matter as well as (**C, F, I**) grey-to-white-matter elasticity ratios *K_g_/K_w_* for (**A-C**) transverse, (**D-F**) horizontal and (**G-I**) sagittal sections. The mean of all animals’ median *K* and the *K_g_/K_w_* ratios (i.e. median grey matter elasticity/median white matter elasticity) are represented by coloured dots; the colour represents the apparent elastic modulus or the ratio, respectively, as shown in the colour bars. The colour of the shaded areas between data points is interpolated for better visualisation. While the elasticity of both grey and white matter increases with increasing force and measurement speed, their ratio *K_g_/K_w_* drops. The number of animals (N) and AFM measurements (n) as well as the individual median grey and white matter elasticity values and *K_g_/K_w_* ratio of each animal, respectively, are provided in Tables S1 and S2. In total, 27,940 measurements contributed to this dataset. For the projection of these plots showing measured elasticity versus force or measured elasticity versus speed, see Figures S3 and S4, respectively.

First, we confirmed that the AFM setup accurately achieves the setpoint forces and speeds over this range (Figure S1). Measurements that deviated from the target values by more than 10% (e.g. due to exceeding the AFM’s z-movement range and thus not reaching the force setpoint) were excluded. Next, we assessed whether AFM measurement themselves altered mechanical tissue properties if conducted with the forces and speeds in this range. We took repeated measurements at the same location and compared the resulting tissue elasticity from consecutive measurements. As the *K* values did not differ to a biologically relevant degree (median value changes by −2%, +4% and +5% in the transverse, horizontal and sagittal plane, respectively; all three planes: p <0.0001) (Figure S2), we averaged data from repeated measurements for further analysis.

We found that in both tissue compartments (i.e. grey and white matter) in all sectioning planes, except in transversely cut white matter, an increase in either force or speed generally led to an increase in the measured apparent elastic modulus. Increases at the lower end of the examined parameter space for both force and speed, from 30 to ∼300 nN and from 20 to 100 µm/s, had a comparably greater effect than further increases of either parameter, indicating a non-linear viscoelastic response (Figure 2; Figures S3 and S4). In transversely cut white matter, however, speed changes had little effect on *K* (Figure 2B; Figures S3B and S4B).

In all anatomical planes, grey and white matter were differentially affected by the tested measurement parameters. As force and speed were increased, grey matter stiffened relatively less than white matter, and thus the grey-to-white matter elasticity ratio *K_g_/K_w_* predominantly decreased with increasing force and speed (Figure 2C, F, I). All mean *K_g_/K_w_* ratios were greater than 1, indicating that across the anatomical planes and parameter combinations tested here, grey matter was stiffer than white matter. However, the smallest mean *K_g_/K_w_* ratio was 1.02 in the sagittal plane, thus approaching 1 and indicating that the measured grey and white matter tissue elasticity were nearly identical. In the sagittal plane, the *K_g_/K_w_* ratio fell below 1 in a single animal, when forces of ≥ 600 nN were applied at speeds of ≥ 400 µm/s, indicating that the measured tissue elasticity of white matter exceeded that of grey matter with these parameter combinations (Figure 3; Table S2).

**Figure 3:**
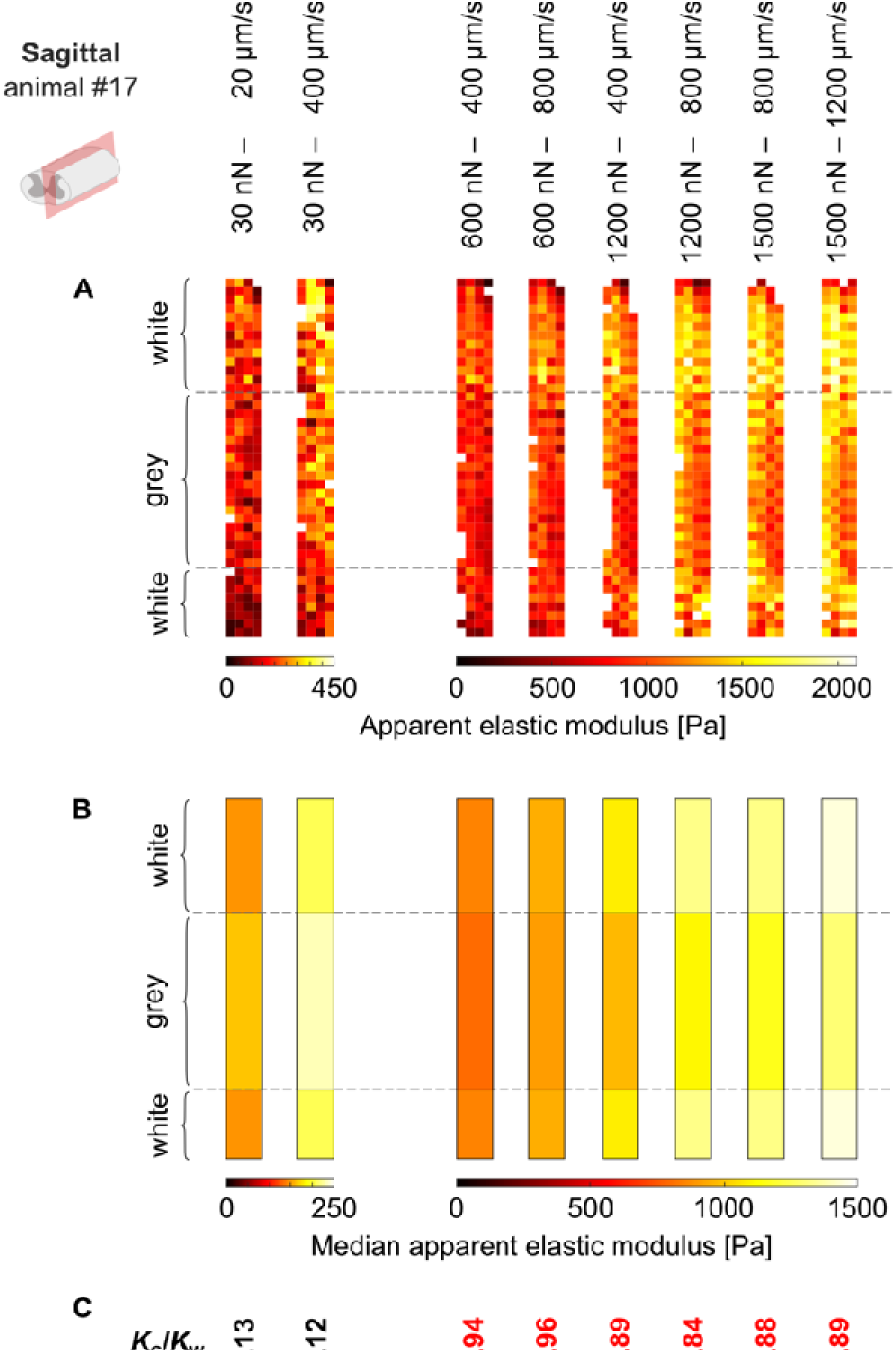
*K_g_/K_w_* ratio can fall below 1 at high forces and speeds. In one animal (#17) measured in the sagittal plane, *K_g_/K*_w_ ratios were <1 for forces of ≥600 nN at speeds ≥400 µm/s. (**A**) Individual AFM measurements and (**B**) median grey and white matter *K* values for all parameter combinations tested in this animal. Parameter combinations are annotated at the top of the figure and relate to all panels. (**A**) Each measurement or (**B**) median grey and white matter *K* values are represented by a colour-coded pixel or rectangle, respectively, illustrating the local apparent elastic modulus as shown in the colour bars. For clarity, data acquired with 30 nN force are displayed with separate colour bars. Dashed black lines: grey-white-matter boundary. (**C**) *K_g_/K_w_* ratio in this animal for every parameter combination. Ratios below 1 are indicated in red.

### K_g_/K_w_ differs across anatomical planes and is highest in transverse sections

While grey matter elasticity was not significantly different across the three anatomical planes (Figure S5), we found anisotropy in white matter elasticity for certain parameter combinations (Figure S6). Despite not being statistically significant in all cases, white matter was consistently softest in the transverse plane (where axon tracts were cut predominantly perpendicular to their long axis and likely relaxed) and stiffer in the horizontal and sagittal planes (where tissue was cut parallel to most axon tracts). Grey matter, on the other hand, displayed a general but not statistically significant trend to be stiffest in the transverse plane. As a result, *K_g_/K_w_* was significantly higher in the transverse plane compared to either of the other two longitudinal planes for all parameter combinations assessed. Between the horizontal and the sagittal planes, *K_g_/K_w_* did not differ significantly (Figure S7).

### Tissue elasticity decreases with time after death, whereas K_g_/K_w_ increases and then plateaus

From here, all experiments were conducted on transverse spinal cord sections with a setpoint force of 30 nN at a speed of 20 µm/s. To investigate the effect of post-mortem time on tissue elasticity, we conducted AFM measurements between 1:30 h and 11h post-mortem (Table 2). Time points before 1:30 h post-mortem could not be assessed due to the time required for the complex sample preparation and AFM setup procedure.

We repeatedly measured the same region of interest every 30 to 60 mins at ∼32.5°C. We confirmed that multiple repeated AFM measurements did not alter the mechanical properties of the tissue by exploiting the mirror-symmetrical structure and mechanics of the spinal cord across the midline. We compared the elasticity changes occurring in one half of a spinal cord section which had been continuously remeasured (seven times in total) with those in the other half of the same section where measurements were conducted only once at the start and once at the end of the experiment and confirmed that the changes in both halves were not significantly different (Figure S8).

Grey matter tissue elasticity was initially relatively stable until ∼3 hours post-mortem. After ∼3 hours, it started declining until ∼6 hours post-mortem, when grey matter elasticity reached another plateau of approximately two thirds of its starting values. In the white matter, by contrast, a pronounced elasticity decrease was apparent immediately from the start of the observation period until ∼6 hours post-mortem, before white matter tissue elasticity plateaued at approximately 50% of its starting values. As a result, the grey-to-white-matter elasticity ratio *K_g_/K_w_* increased slightly until ∼6 hours post-mortem before it levelled out (Figure 4). However, the dynamics of the *K_g_/K_w_* ratio were overall relatively variable across different animals.

**Figure 4:**
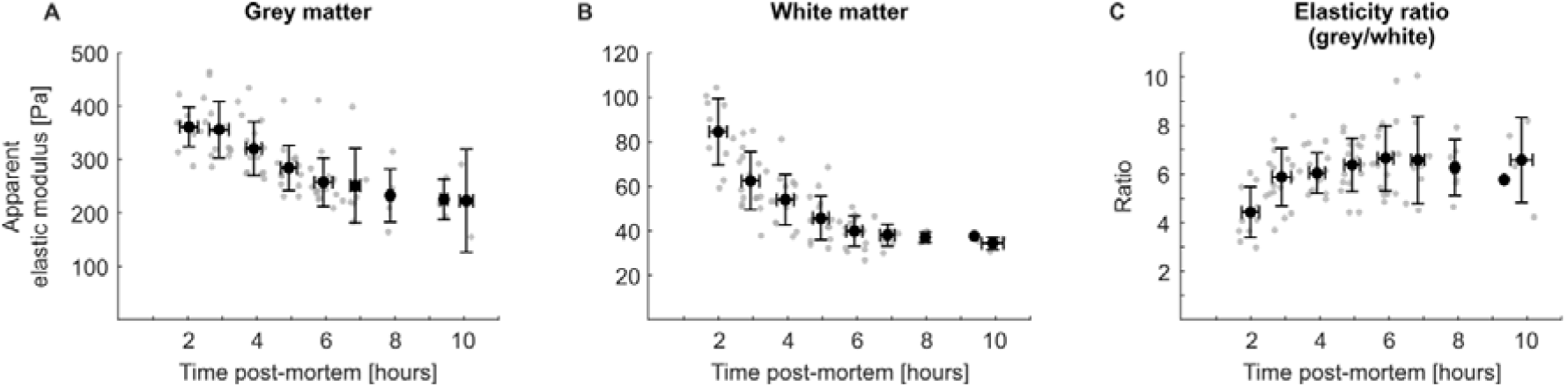
Grey and white matter tissue elasticity decline with time after death, whereas grey-to-white-matter elasticity ratio increases. Transverse spinal cord sections of eleven rats were used to record AFM maps 6 – 13 times every ∼30 – 60 mins. Data are shown separately for (**A**) grey matter, (**B**) white matter and (**C**) the ratio of grey-to-white-matter elasticity. While grey matter elasticity only started dropping after ∼3 hours post-mortem, white matter elasticity declined from the beginning. Both tissue compartments reached a plateau after ∼6 hours post-mortem. As white matter elasticity declined faster than grey matter elasticity, *K_g_/K_w_* increased until ∼6 hours post-mortem. Grey dots: (**A, B**) median *K* over median time post-mortem per map (grey matter: 28 – 56 measurements/map/animal, in total 3,591 measurements; white matter: 36 – 95 measurements/map/animal, in total 5,300 measurements) or (**C**) the ratio of median grey to median white matter elasticity over the mean of the median times post-mortem at which the map was acquired. Black dots with error bars denote mean ± SD for one-hour bins, starting at 1:30 h post-mortem, of data shown in grey. Data were acquired with 30 nN force at 20 µm/s speed. Maps which were not completed in time were excluded from data analysis, resulting in 6 – 11 usable maps per animal.

### Temperature affects absolute tissue elasticity but not K_g_/K_w_

Finally, we investigated the effect of temperature on spinal cord tissue elasticity in the range of 38°C to 20°C, representing physiological rat body temperature and typical room temperature, respectively. Grey and white matter tissue elasticity both increased as temperature decreased (Figure 5A, B; Table 3). As grey and white matter elasticity changed at an almost identical rate (grey matter: 3.9% per 1°C; white matter: 3.7% per 1°C; see Table 3), *K_g_/K_w_* was very stable in the examined temperature window and independent of the temperature (Figure 5C).

**Figure 5:**
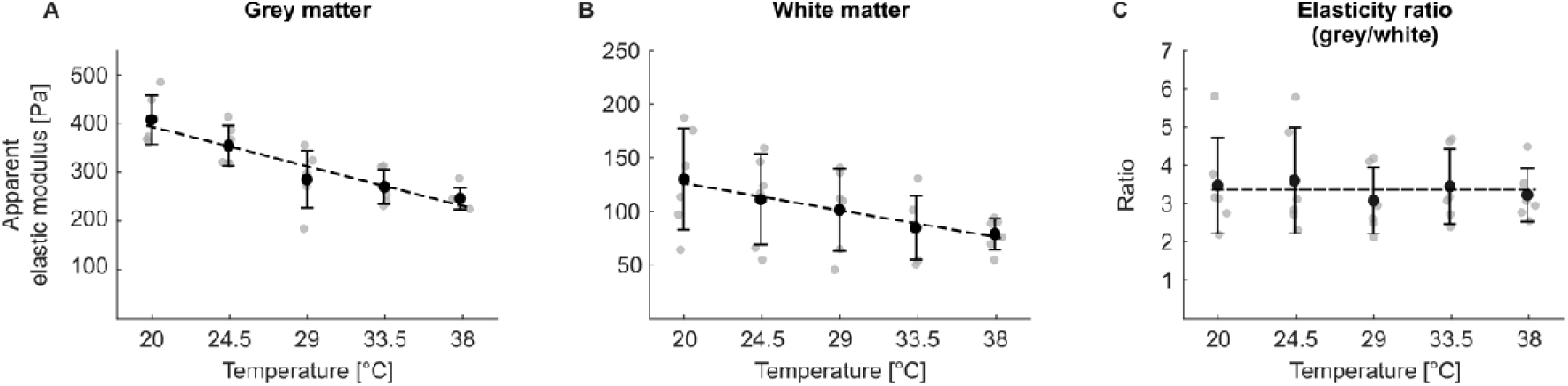
Grey and white matter tissue elasticity decrease with increasing temperature while the ratio of grey-to-white-matter elasticity is unaffected. AFM maps were repeatedly recorded at five temperature levels (20°C, 24.5°C, 29°C, 33.5°C and 38°C). Data are shown for (**A**) grey matter, (**B**) white matter and (**C**) the ratio of grey-to-white-matter elasticity. Grey dots: (**A, B**) median *K* per animal over median measured temperature (grey matter: 35 – 48 measurements/temperature level/animal, in total 1,308 measurements; white matter: 44 – 62 measurements/temperature level/animal, in total 1,680 measurements) or (**C**) the ratio of median grey to median white matter elasticity per animal over the mean of the median measured temperatures for grey and white matter. Black dots with error bars: mean ± SD of data shown in grey, binned for each temperature level. The SD along the temperature axis was very small (< 0.33°C) and is not shown for clarity. Black dashed lines: (**A, B**) Linear regressions; (**C**) horizontal line. The grey-to-white-matter elasticity ratio is not dependent on temperature. Regression analysis details are provided in Table 3. Data were acquired with 30 nN force at 20 µm/s speed.

**Table 3:**
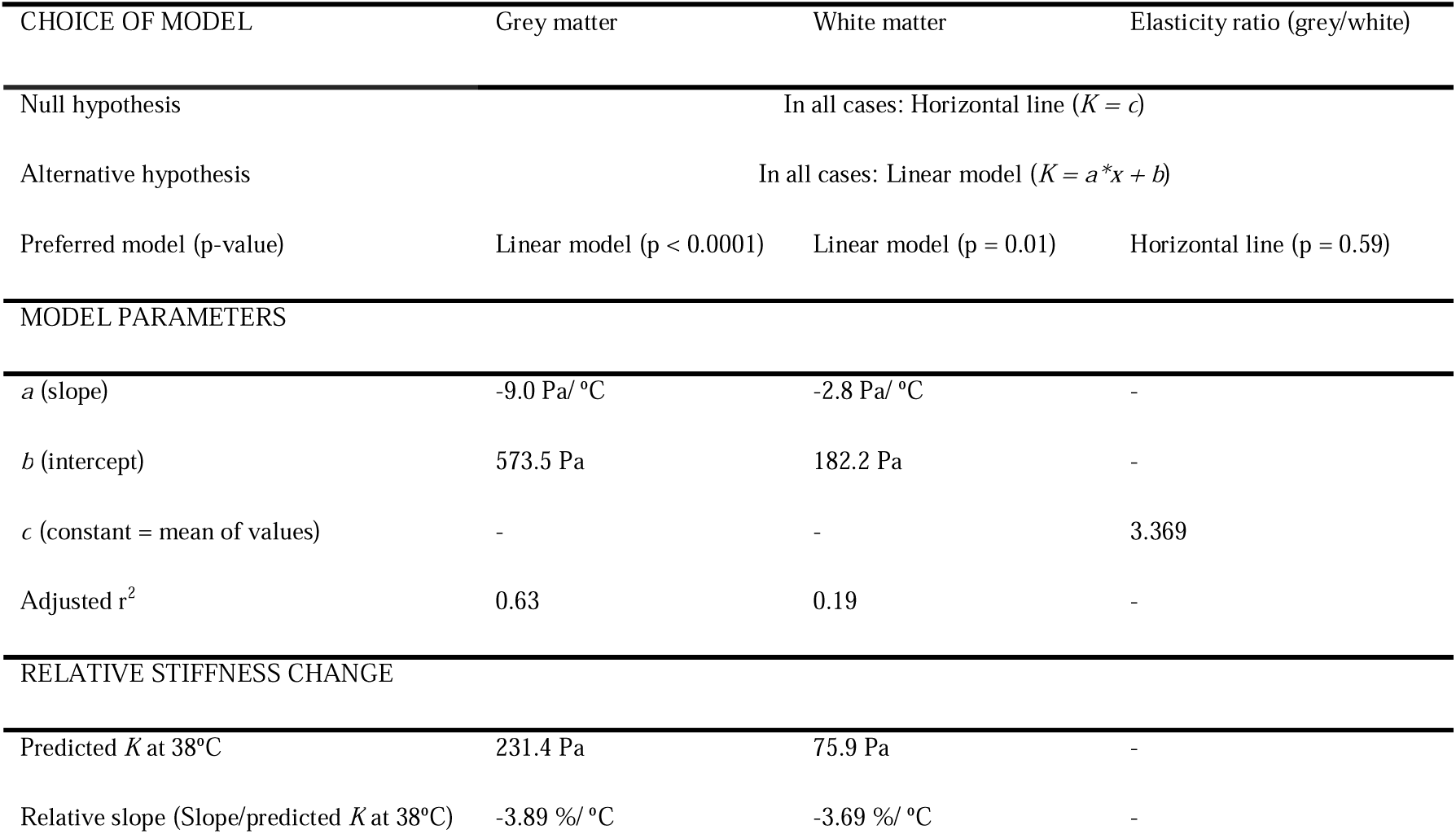
Regression analysis of tissue elasticity *K* [Pa] as a function of temperature *x* [°C]. This table relates to Figure 5. Regression analysis was conducted on all animals’ median *K* values (see Figure 5, grey dots). A linear regression model was compared to a horizontal line, i.e. the assumption that temperature did not affect tissue elasticity, with an extra-sum-of-squares F Test. Despite grey and white matter elasticity in themselves being temperature-dependent, the grey-to-white-matter elasticity ratio was not affected by temperature, as the relative elasticity changes in grey and white matter (normalised to the predicted *K* at body temperature, i.e. 38°C) were similar.

| CHOICE OF MODEL | Grey matter | White matter | Elasticity ratio (grey/white) |
| --- | --- | --- | --- |
| Null hypothesis | In all cases: Horizontal line ($K = c$) | | |
| Alternative hypothesis | In all cases: Linear model ($K = a*x + b$) | | |
| Preferred model (p-value) | Linear model (p < 0.0001) | Linear model (p = 0.01) | Horizontal line (p = 0.59) |
| MODEL PARAMETERS |  |  |  |
| $a$ (slope) | -9.0 Pa/ °C | -2.8 Pa/ °C | - |
| $b$ (intercept) | 573.5 Pa | 182.2 Pa | - |
| $c$ (constant = mean of values) | - | - | 3.369 |
| Adjusted $r^2$ | 0.63 | 0.19 | - |
| RELATIVE STIFFNESS CHANGE |  |  |  |
| Predicted $K$ at 38°C | 231.4 Pa | 75.9 Pa | - |
| Relative slope (Slope/predicted $K$ at 38°C) | -3.89 %/ °C | -3.69 %/ °C | - |

## Discussion

We have shown that the measured elasticity of spinal cord tissue strongly depends on the force and speed at which measurements are conducted. We confirmed that the AFM measurements themselves did not alter measured spinal cord elasticity to a biologically relevant degree within the explored parameter space (Figure S2). To investigate the effect of directionality, we conducted our measurements in all three anatomical planes (transverse, horizontal and sagittal; see Figure 1 F, G, H). As force and speed increased, spinal cord stiffened in all tissue sections, in agreement with previous compressive tests in the CNS (10, 14, 15, 20, 21, 30). However, we found region-specific differences in the tissue’s strain stiffening behaviour: grey matter stiffened relatively less than white matter with increasing forces and speeds, so that the ratio of grey-to-white-matter elasticity *K_g_/K_w_* decreased (Figure 2; Figure S3). In one animal, *K_g_/K_w_* fell below 1 for high forces and speeds (Figure 3), but generally grey matter was stiffer than white matter within the explored parameter space. The distinct strain stiffening behaviour of grey and white matter elasticity could be related to their different microarchitecture. Axons contain large amounts of intermediate filaments (neurofilaments), which show a very strong strain stiffening behaviour if compared to other cytoskeletal components (31), potentially explaining why white matter stiffens relatively more than grey matter when exposed to larger strains.

Our experimental settings extended over two orders of magnitude, with forces ranging from 30 to 1500 nN and speeds ranging from 20 to 1200 µm/s. With indenters of radius *R* = 44.65 µm, this resulted in indentation depths of 4 – 40 µm and indentation durations of 8 – 1278 ms in 95% of all measurements. Even though AFM settings do not directly translate into strains and strain rates, we estimate the nominal strain *ε* = Δ*l*/*l*_0_ (Eq. 1), where Δ*l* is the length change, *l*_0_ the original length, *δ* the indentation depth (4 – 40 µm) and *h* the sample height (999 µm), to be between ∼0.4 – 4% and the strain rate 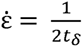 (Eq. 2), where *t_δ_* is the indentation duration (8 – 1278 ms), to be approximately 0.4 – 60 Hz. Of note, local strains directly below the indenter will have been higher as the strain field is not uniform.

Both nanoindentation testing and MRE studies often find white matter to be stiffer than grey matter (Table 4). In nanoindentation measurements, indentations of between ∼5 to 10% of the sample height are commonly reported (14, 15, 32), sometimes even up to 20% (33), at speeds similar to the lower end of our parameter space (5 – 100 µm/s (14–16, 34)). MRE, on the other hand, exerts deformations of about 1 to 40 micrometres in a large sample like a human brain (35–37), resulting in smaller strains then used in this study, but at higher strain rates of about 50 – 100 Hz (35–40). In our study, mean *K_g_/K_w_* values approached 1 for high forces and speeds in the horizontal and sagittal planes, and we even observed *K_g_/K_w_* falling below 1 in one animal (Figure 3). This suggests that a *K_g_/K_w_* < 1 as observed with nanoindentation and MRE could at least partially be the result of the even higher forces or speeds employed by these methods (Table 4). Additionally, the much larger probe size of nanoindenters on the order of 750 – 1000 µm radius (14–16, 32–34) could lead to measuring collective behaviours of axon bundles at larger scales. Also, CNS exhibits both viscoelastic behaviour (because of rearrangements of cells and the extracellular matrix) and poroelastic behaviour (because of fluid flows through the matrix) (41). Differences in water displacement in grey compared to white matter could result in white matter appearing stiffer during shear wave propagation in MRE.

**Table 4:**
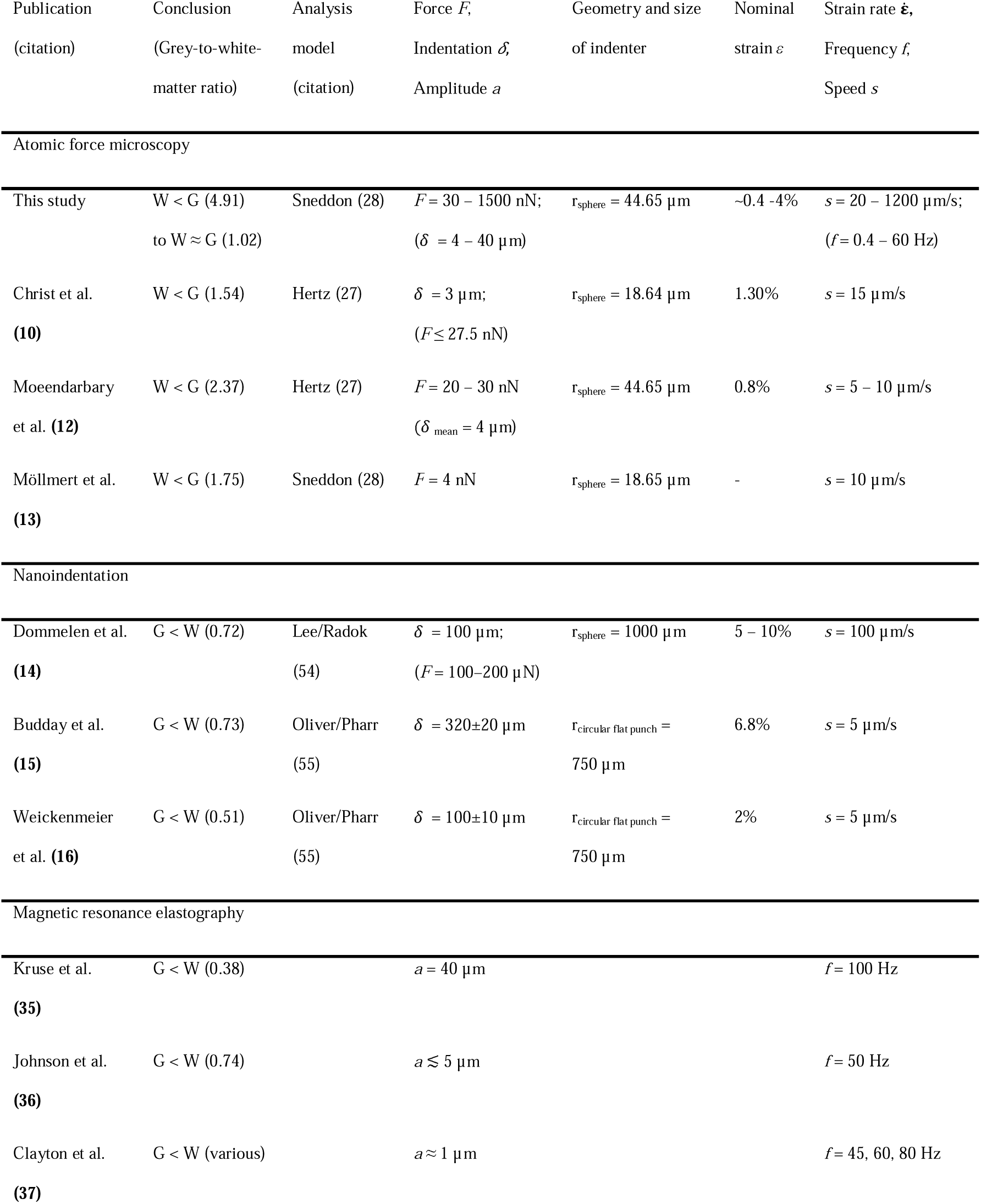
Comparison of controversial data concerning grey and white matter mechanical properties in studies employing different strains and strain rates. Selected publications sorted by methodology. Strain-related parameters: Force *F*, Indentation, Amplitude *a*. The controlled variable is indicated first. If such data were available, the resulting variable is indicated second in brackets. Nominal strain *ε* was calculated based on Eq. 1 where sufficient data were available in the respective publication. Strain rate-related parameters: Frequency *f*, speed *s.* G = grey matter; W = white matter.

*K_g_/K_w_* was significantly higher in the transverse plane (perpendicular to the main direction of axonal fibre tracts) than in either the horizontal or the sagittal planes (both parallel to the main axonal fibre direction) (Figure S7). In both longitudinal planes, *K_g_/K_w_* values were similar and closer to 1. Thus, measurement directionality relative to the predominant orientation of white matter tracts strongly affected *K_g_/K_w_*. We found grey matter to be isotropic with all parameter combinations and white matter to be transversely isotropic with some parameter combinations but anisotropic if comparing longitudinal and transverse directions, which agrees with previous findings from AFM and MRE studies (11, 42). Overall, both grey and white matter elasticity decreased over time post-mortem. Grey matter elasticity was initially stable for ∼3 hours. It then decreased, before stabilising again at ∼6 hours post-mortem at a lower level. In contrast, white matter displayed a pronounced drop of tissue elasticity in the first 6 hours post-mortem. Consequentially, the grey-to-white-matter elasticity ratio increased initially before plateauing after approximately 6 hours post-mortem (Figure 4). Some previous studies reported brain tissue elasticity not to change for up to 5 days post-mortem (15) or to start increasing ∼6 hours after an initial stable phase (22). Tissues in these studies had been sourced from an abattoir, so it is possible that there were delays between death and brain tissue harvest. Furthermore, brains were stored in PBS, which cannot mitigate the complex metabolic changes affecting CNS tissue post-mortem (26). It is therefore possible that the tissues used in these studies may have undergone mechanical changes already before the start of measurements, possibly even reaching a plateau phase as seen in the later time points of our data in both grey and white matter (Figure 4A, B).

MRE studies measuring mechanical brain tissue properties *in vivo* and again within ∼30 – 60 minutes post-mortem *in situ* reported a considerable increase in tissue stiffness (23, 43–45). This has been attributed to cytotoxic oedema and brain swelling inside the spatially confined cranial cavity (23). If the brain was promptly taken out of the skull after death, the measured tissue elasticity was, in contrast, considerably lower than *in vivo*, which might be due to a sudden lack of blood and cerebrospinal fluid pressure (24). The studies agree that after these immediate post-mortem events, the measured tissue stiffness remains constant in the early post-mortem phase until ∼1 to 3 hours post-mortem (23, 24, 44), compatible with our findings in grey matter (Figure 4A). At later time points, tissue stiffness was reduced (43, 44), which further agrees with our findings in both grey and white matter.

We here identified clear differences in the susceptibility of grey and white matter elasticity for the passing of time. Using buffer solutions specifically optimised to ensure spinal cord slice viability and normal electrical activity of grey matter neurons for hours after slice preparation (26) likely helped slowing down the decay of the grey matter. In contrast, severing CNS axons *in vivo* leads to progressive axonal swelling, structural defects of the axonal cytoskeleton and the myelin sheath, and the accumulation of vacuoles within the first six hours (46), potentially explaining the rapid decrease in white matter elasticity post-mortem.

We confirmed temperature to be another important parameter impacting measured tissue elasticity. Tissue elasticity linearly increased as temperature decreased from physiological body temperature (38°C) to room temperature (20°C), in line with previously published results (24, 25, 47, 48) (Figure 5A, B). Here, we found that the relative elasticity increase was similar for grey and white matter, and thus the grey-to-white-matter elasticity ratio was unaffected by temperature (Figure 5C). The liquid-crystal to gel-phase transition of myelin takes place at 63°C (49), so that considerable differences in the mechanical behaviour of grey and white matter would only be expected above this threshold. This rules out that different measurement temperatures in past studies are responsible for the differences in the reported *K_g_/K_w_* values.

## Limitations

While we have attempted to address some major contributors to reported differences in the grey-to-white-matter elasticity ratio, other factors might still play a role. Due to the scarcity of data available for the spinal cord specifically, we have compared our findings to data obtained on CNS tissue in general, i.e. both brain and spinal cord tissue. Both form a functional unit and contain grey and white matter. However, local mechanical differences exist even within grey and white matter (42, 50), so that this division, while practically useful, remains somewhat crude. More work is needed to better understand the spatial heterogeneity in different brain and spinal cord regions and white matter tracts.

The anatomical context in which CNS tissue is examined is also important. While MRE allows for in vivo measurements, many other methods require post-mortem tissue. As white matter is under tension (51, 52), and this tension may be released during sample preparation, *ex situ*/*in vitro* measurements may not fully recapitulate *in vivo*/*in situ* mechanics. Furthermore, it is important to note that our AFM measurements constitute compressive testing and that measurements in tension or in shear could yield different results.

The fact that different methods use different models for data analysis and report different outcome variables poses a general problem for literature comparisons in the field and likely also contributes to inconsistent findings. The standard models to analyse AFM data are the Hertz and the Sneddon model. The Hertz model (27) is simpler and considered appropriate for low indentation depth relative to the indenter size, whereas the Sneddon model (28) has to be solved numerically but is accurate also at higher indentation depths. Here, we have used the Sneddon model for experiments examining the effect of different force-speed combinations, due to the greater indentation depths, and the Hertz model for all other experiments. Both models assume the sample to be isotropic, homogeneous and linearly elastic. None of these conditions are met entirely by biological samples in the real world, but we designed our experiments to account for these violations. Samples were taken from specified anatomical locations and orientations to account for the lack of isotropy, and measurements were conducted in a spatially resolved way with a specified indenter size to account for the lack of homogeneity. We were working in a low-strain regime (*F* = 30 nN with the given indenter size), where the sample can be assumed close to linearly elastic. Despite exerting much higher forces in parts of this study, the Sneddon model still fit the force-indentation data curves well (Figure 1 B-E). We also confirmed that the apparent reduced elastic modulus *K*, which we obtained with our analysis, was highly correlated with another commonly used – model-independent – measure, the stiffness *k* (unit: N/m), which we obtained by fitting a linear function to the force-indentation data in the range of 90 – 100% of the exerted force (Figure S9; Pearson’s *r* = 0.95). However, other physical models can also take viscoelastic or poroviscoelastic effects into consideration, which would affect absolute measured values and possibly also *K_g_/K_w_*. Conducting creep or oscillatory measurements with AFM to assess viscoelastic effects in a spatially resolved way would be a great future asset for the community.

## Conclusions

Force, speed, and post-mortem interval all influenced the grey-to-white-matter elasticity ratio *K_g_/K_w_*. Our results therefore reconcile seemingly contradictory findings in the literature concerning the relative elasticity of grey and white matter obtained with methods employing very different strain and strain rate regimes. Given the tissue compartment-specific dependence of elasticity on these parameters, it is important to tune the choice of method to the question being asked. If the mechanical environment experienced by cells *in vivo* is in focus, small strains and strain rates should be used. If data is to be acquired for modelling of high impact scenarios such as traffic accidents, high strains and strain rates should be chosen in order to learn how the tissue behaves under these conditions.

Furthermore, high-quality and timely sample preparation is paramount for reliable measurements of CNS tissue elasticity, in particular if white matter is concerned. Attention also must be paid to the method chosen for euthanasia, which can impact mechanical tissue properties (53), and immediate supply of (i.e. perfusion with) a physiological buffer, such as artificial cerebrospinal fluid solutions (26), must be ensured. Careful consideration of sample preparation techniques and selection of measurement parameters, which mimic the relevant *in vivo* situation more closely, will lead to more impactful results, as well as better standardisation and comparability of results across techniques and laboratories.

Our findings therefore suggest that the contradictory findings in the field might at least partly be due to different strain and strain rate regimes employed by different measurement methods, as well as testing being conducted along different anatomical axes, and illustrate the complex non-linear nature of CNS tissue mechanics. Future research should investigate if further parameters such as probe size and poroviscoelastic properties of CNS tissue also contribute to the apparently contradictory *K_g_/K_w_* values reported in the literature.

## Author contributions

JMB, EK and KF designed the study; JMB performed the experiments and analysed the data; AKW provided technical support and contributed software; JMB and KF wrote the manuscript, with contributions from all co-authors.

## Acknowledgements

The authors thank the animal facility staff for their excellent work and the research groups which donated animals to this project. This work was supported by the Wellcome Trust (PhD fellowship to JMB), the European Research Council (Consolidator Award 772426 MECHEMGUI and Synergy Grant 101118729 UNFOLD to KF), the German Research Foundation (DFG) (projects in 460333672 CRC1540 EBM and 270949263 GRK2162 to KF), and the Alexander von Humboldt Foundation (Alexander von Humboldt Professorship to KF).

## Supporting material

**Figure S1:**
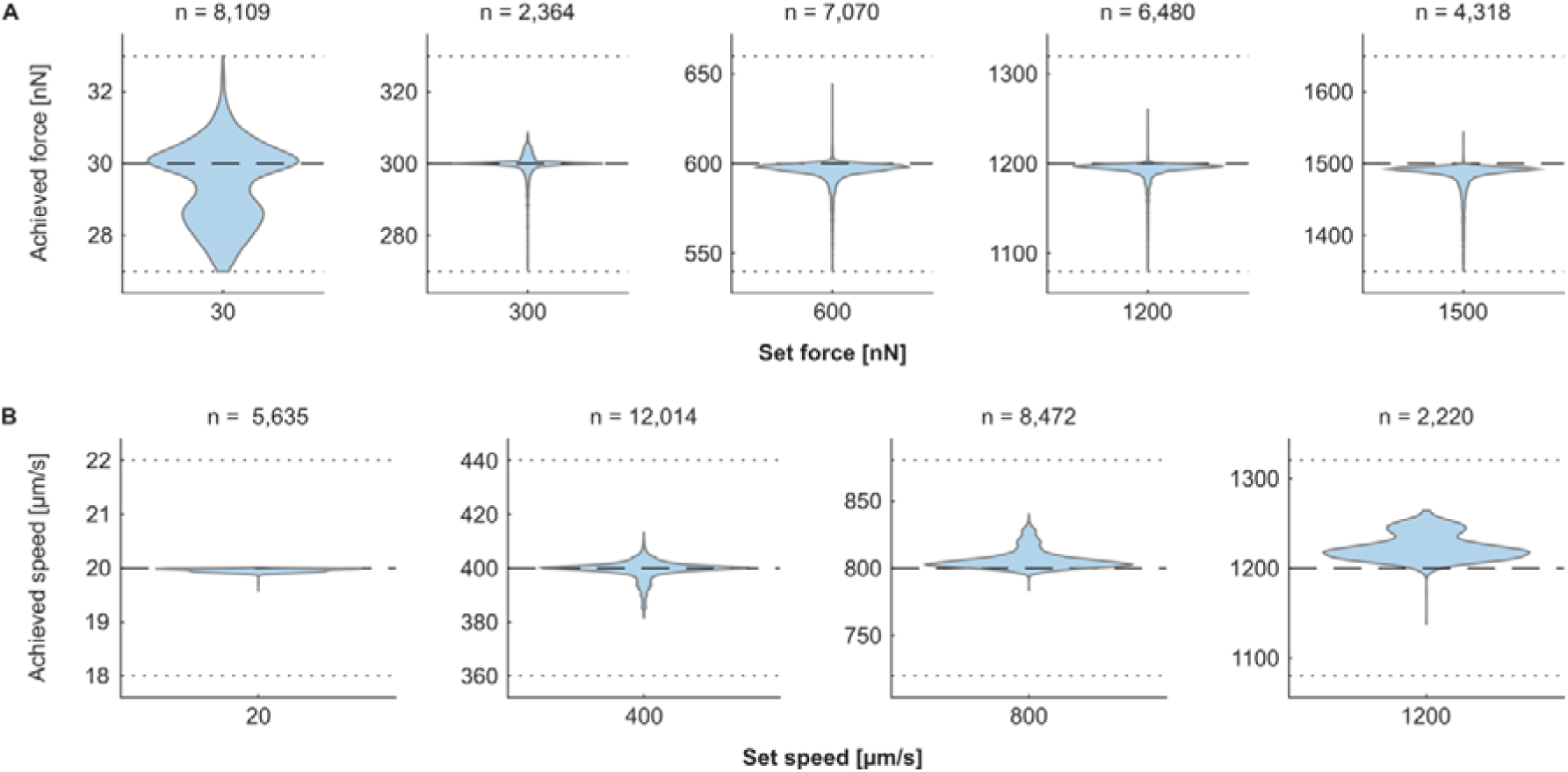
AFM accurately achieves setpoint forces and speeds across a wide range. (**A)** Actually achieved forces and (**B)** speeds plotted against setpoint (= target) forces and speeds. The target forces and speeds are indicated by dashed lines, ±10% of target forces and speeds are indicated by dotted lines. The number of measurements analysed is indicated above each plot. Measurements for which the achieved force deviated by more than 10% from the target force were excluded from further data analysis. No achieved speed exceeded the target speeds by more than ±10%.

**Figure S2:**
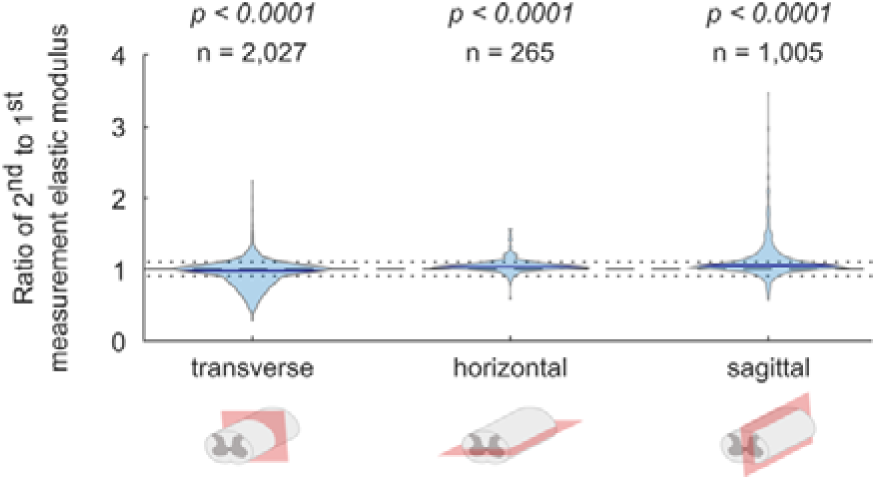
Measurements conducted in the chosen parameter space do not alter the tissue’s apparent elastic modulus to a relevant degree. Data relate to Figure 2. To assess whether AFM measurements with the highest forces and speeds employed there mechanically damage the tissue, AFM measurements were taken with 30 force-speed combinations at the same location and were then repeated once. Apparent elastic moduli of the second measurements were compared to the first ones. Data shown are pooled for each anatomical plane and were compared against a value of 1 with a two-tailed one-sample t-test. The number of measurements per plane (*n*) and the p-values are indicated in the figure. The distribution medians were close to 1 (transverse: 0.981; horizontal: 1.037; sagittal: 1.053; all indicated with blue lines), indicating that no experimentally meaningful mechanical alterations had occurred. The dashed line indicates a ratio of 1, the dotted lines denote 1±10%.

**Figure S3:**
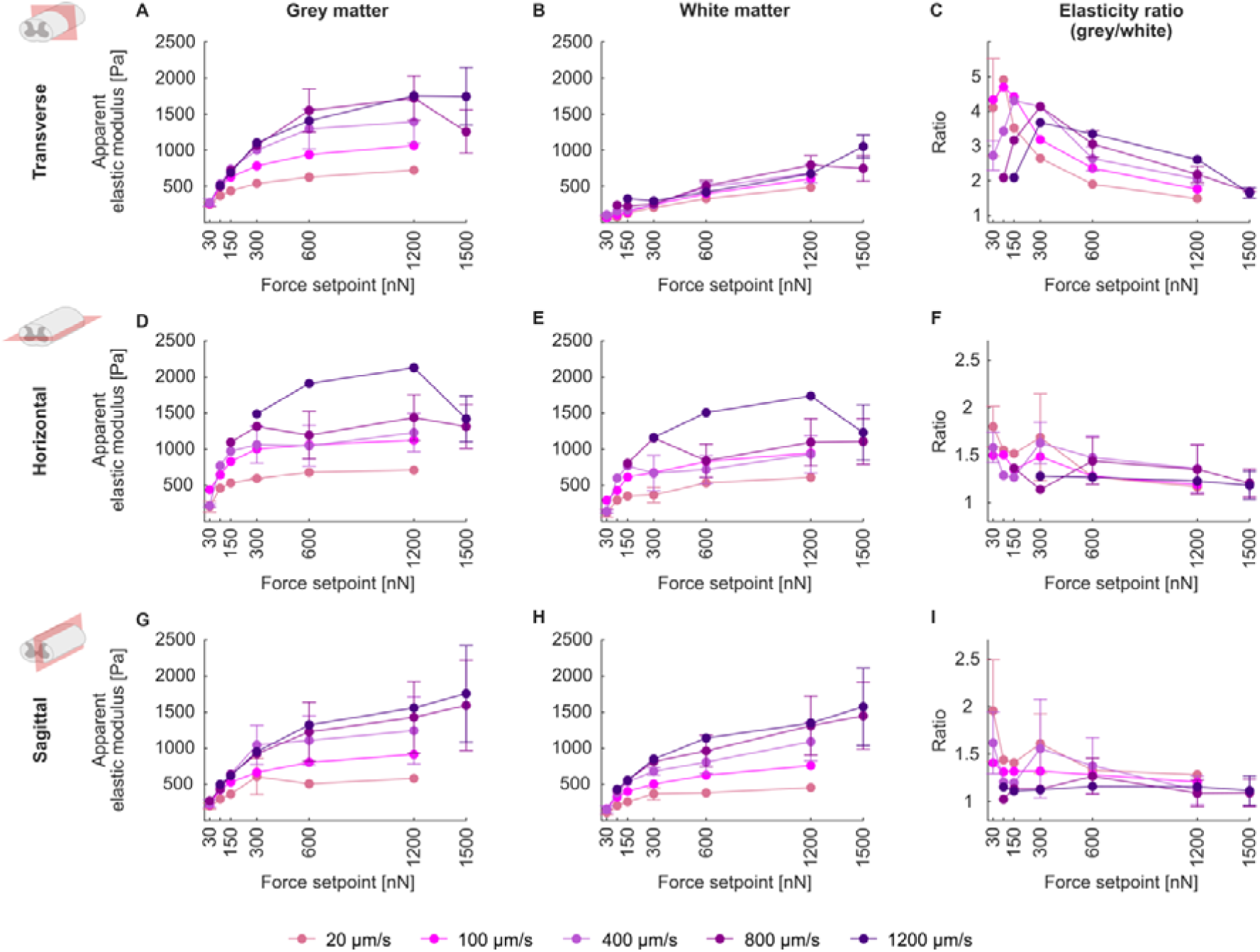
Increasing the force setpoint generally increases grey and white matter elasticity and decreases the grey-to-white-matter elasticity ratio. Projections of data shown in Figure 2, showing measured elasticity versus force. Dots and error bars represent means and standard deviations of all animals’ median *K* (**A, B, D, E, G, H**) or *K_g_/K_w_* ratios (**C, F, I**). Where error bars are missing, N = 1 animal. For details about animal numbers, measurement numbers and median elasticity values and ratios from individual animals, please refer to Tables S1 and S2. Data are colour-coded according to the setpoint speed at which they were acquired; data acquired at the same speed are connected by lines.

**Figure S4:**
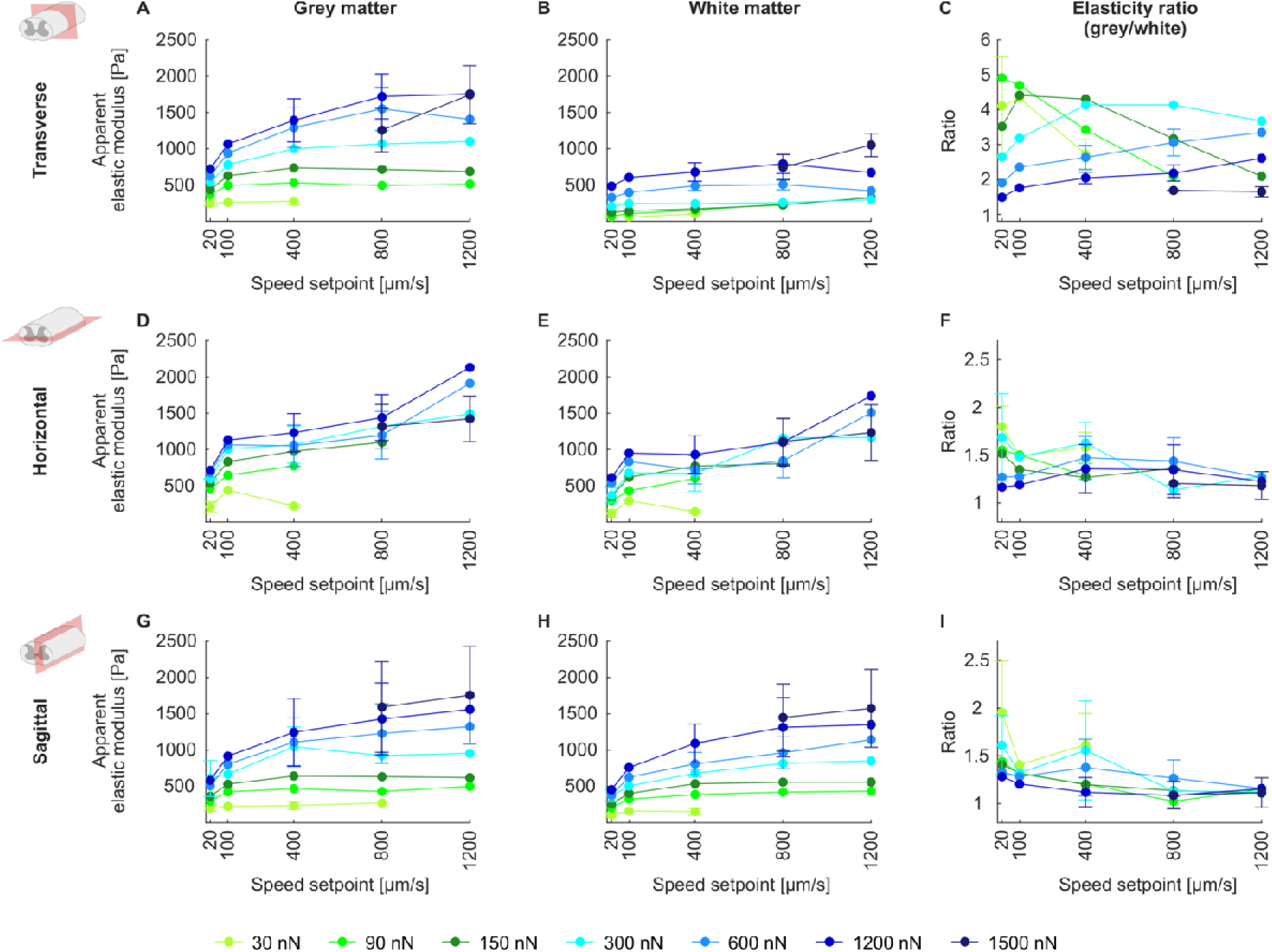
Increasing the speed setpoint generally increases grey and white matter elasticity and decreases the grey-to-white-matter elasticity ratio. Projections of data shown in Figure 2, showing measured elasticity versus speed. Dots and error bars represent means and standard deviations of all animals’ median *K* (**A, B, D, E, G, H**) or *K_g_/K_w_* ratios (**C, F, I**). Where error bars are missing, N = 1 animal. For details about animal numbers, measurement numbers and median elasticity values and ratios from individual animals, please refer to Tables S1 and S2. Data are colour-coded according to the setpoint force at which they were acquired; data acquired with the same force are connected by lines.

**Figure S5:**
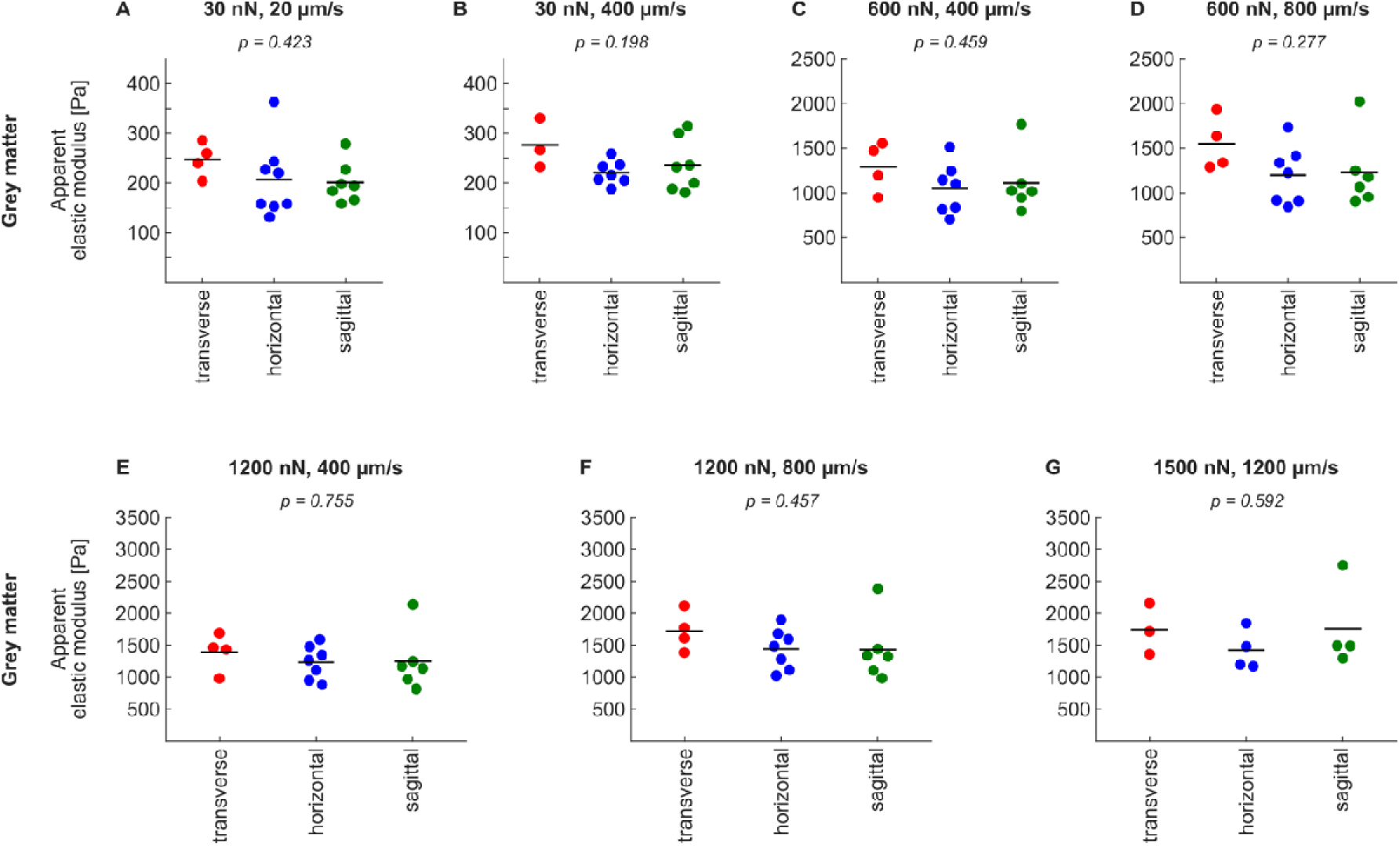
Grey matter elasticity does not significantly differ across the three anatomical planes. Data relate to Figure 2. (**A-G**) Comparison of elasticity values obtained with the same parameter combinations in the three anatomical planes. Dots represent individual animals’ median *K*, bars their means. Anatomical planes were compared with ANOVA. None of the comparisons were significant (p > 0.05 in all cases). Parameter combinations were included in the analysis if data was available for all three anatomical planes and if N ≥ 3 animals/plane (N = 3 – 8 animals/plane). Data shown here are included in Table S1.

**Figure S6:**
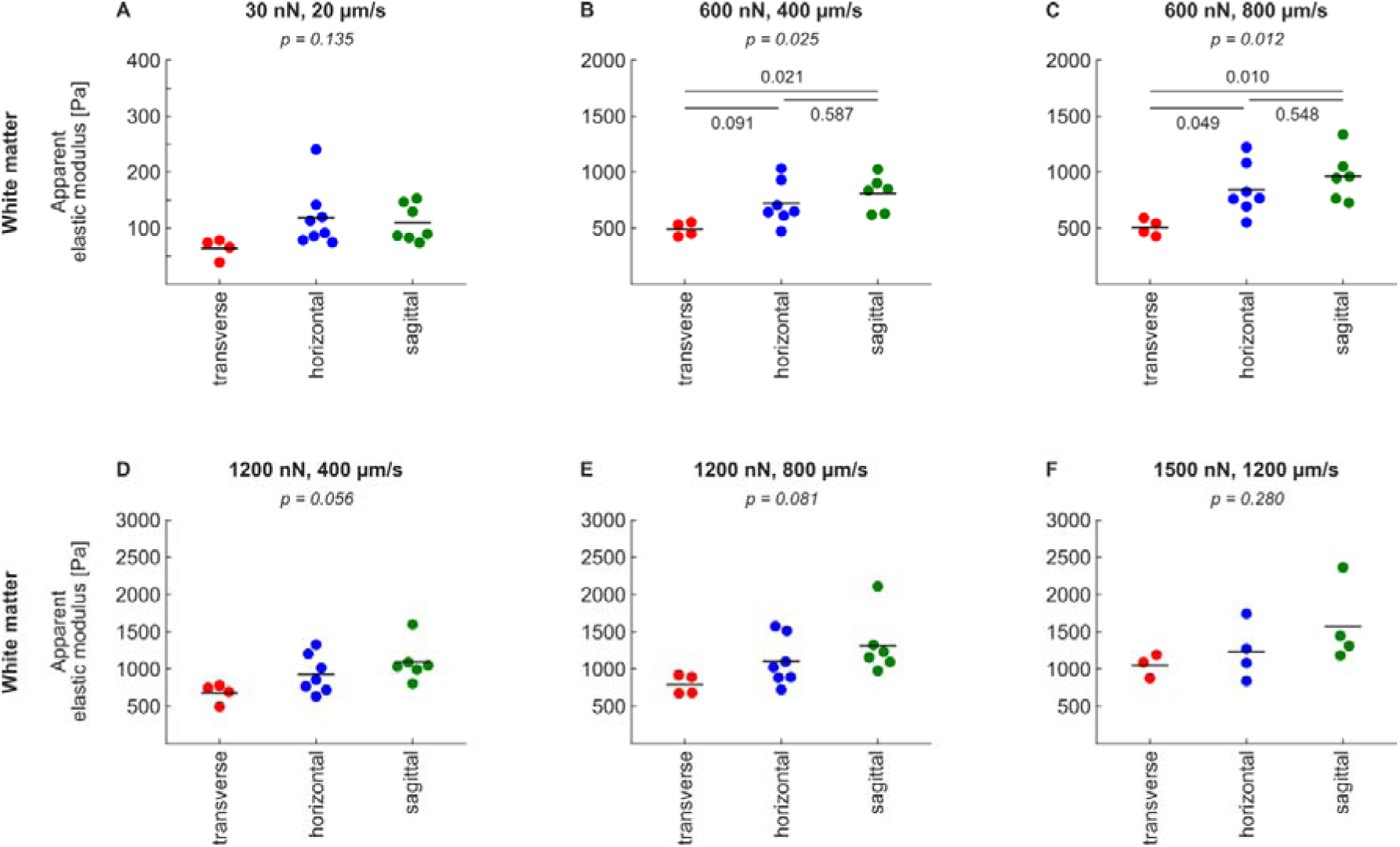
White matter elasticity is lowest in the transverse plane. Data relate to Figure 2. (**A-F**) Comparison of elasticity values obtained with the same parameter combinations in the three anatomical planes. Dots represent individual animals’ median *K*, bars their means. Anatomical planes were compared with ANOVA (p-values in italics at the top of each panel). Where ANOVA yielded p ≤ 0.05 (**B, C**), Tukey’s multiple comparisons test was used to compare individual planes (comparisons indicated by bars with adjacent p-values). Parameter combinations were included in the analysis if data was available for all three anatomical planes and if N ≥ 3 animals/plane (N = 3 – 8 animals/plane). Data shown here are included in Table S1.

**Figure S7:**
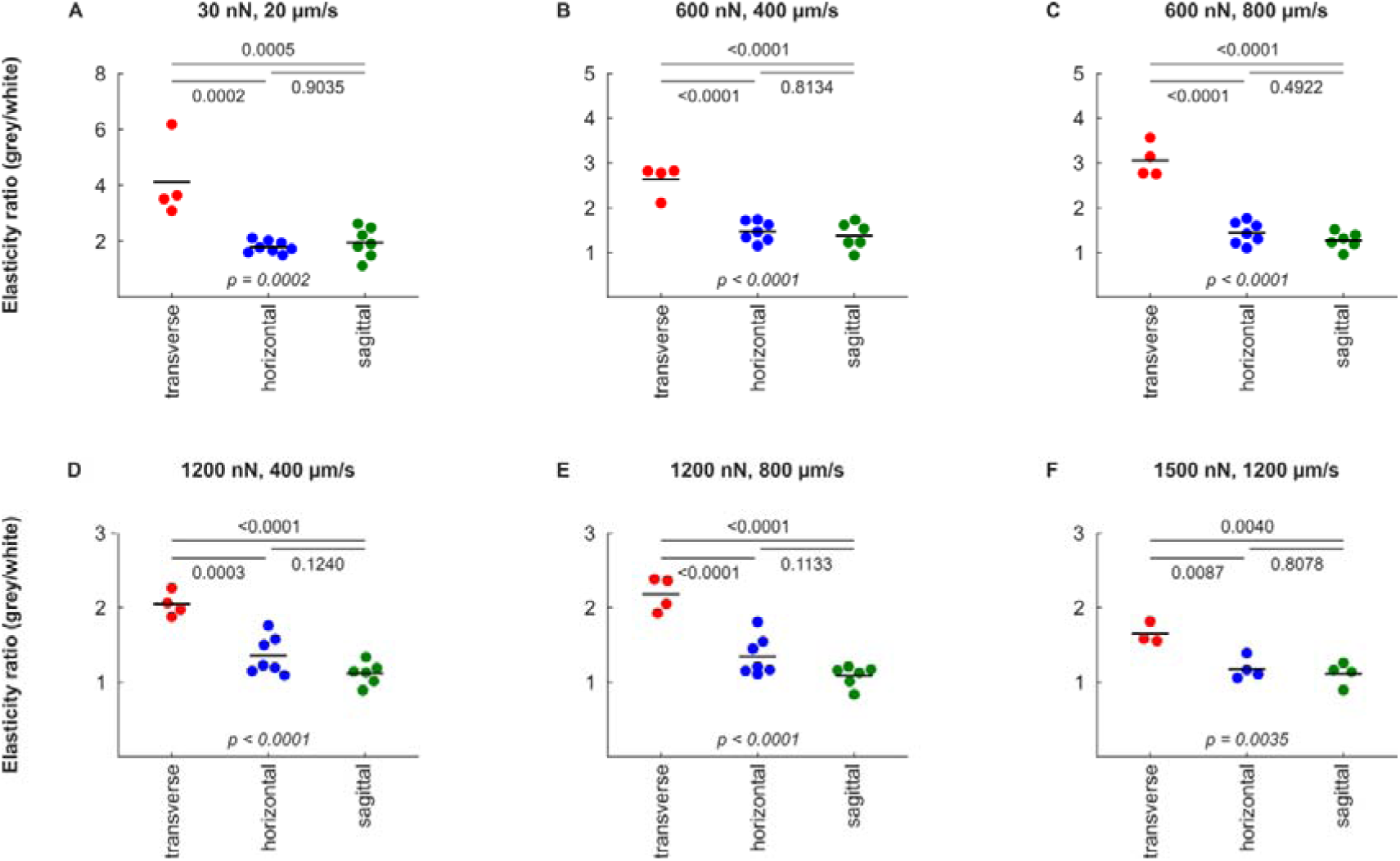
Measurement directionality significantly affects the *K_g_/K_w_* ratio. Data relate to Figure 2. (**A-F**) Comparison of *K_g_/K_w_* ratios obtained with the same parameter combinations in the three anatomical planes. Dots represent individual animals’ *K_g_/K_w_* ratios, bars their means. Anatomical planes were compared with ANOVA (p-values in italics at the bottom of each panel; all significant), followed by Tukey’s multiple comparisons test (indicated by bars with adjacent p-values). In all cases, *K_g_/K_w_* values were significantly higher in the transverse plane than in either the horizontal or sagittal planes. In all cases, *K_g_/K_w_* ratios obtained in the horizontal and the sagittal plane did not significantly differ. Parameter combinations were included in the analysis if data was available for all three anatomical planes and if N ≥ 3 animals/plane (N = 3 – 8 animals/plane). Data shown here are included in Table S2.

**Figure S8:**
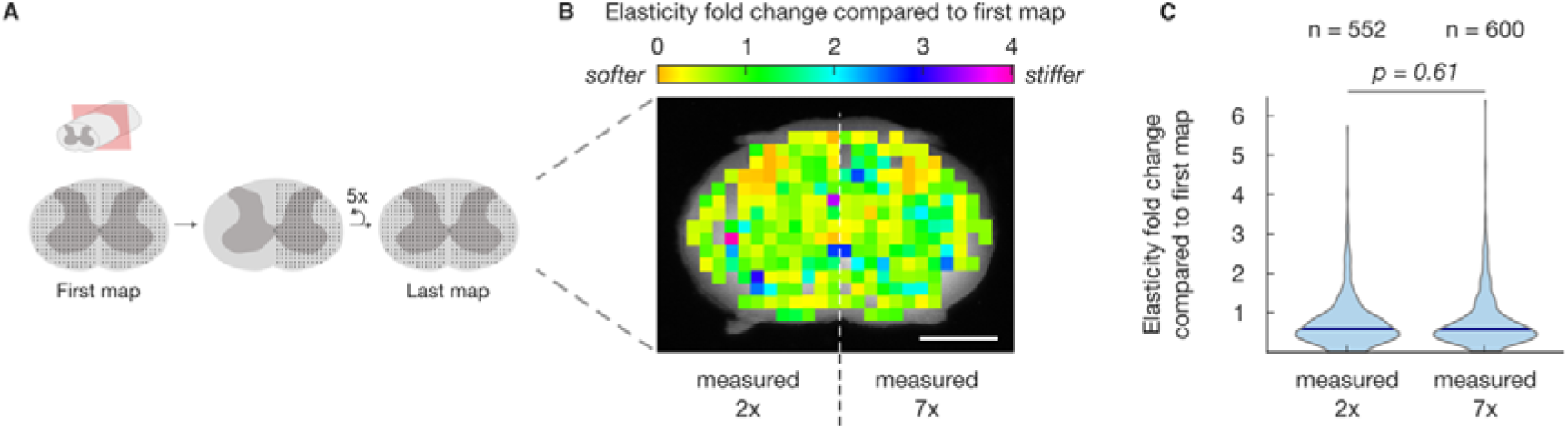
Multiple repeated low-force-low-speed measurements do not alter the apparent elastic modulus of spinal cord tissue. (**A**) Initially, an AFM map was measured on each half of a spinal cord cross-section. Subsequently, only one of the two grids was measured five more times every ∼30 or 60 minutes. Finally, both sides were measured again. The initial maps were acquired between 1:38 h and 3:51 h (mean: 2:49 h) post-mortem, measurements in the last maps between 8:31 h and 11:06 h (mean: 9:45 h) post-mortem. (**B**) Relative elasticity changes between the first and last AFM map in both sides of the spinal cord cross-section in a representative animal. The heatmap illustrates midline symmetry (midline annotated with a dashed line). The left side was measured twice, whereas the right side of the spinal cord was measured seven times within the same time interval. (**C**) Distribution of elasticity changes between the first and last AFM maps in areas measured twice and areas measured 7 times (97 – 126 measurements per map per animal, N = 5 animals). Both conditions were compared with a two-tailed unpaired t-test after log-transformation (p = 0.61). The five additional AFM elasticity mapping experiments did not significantly change the elasticity of the tissue. Median for “measured 2x” is 0.595; median for “measured 7x” is 0.582. Scale bar = 1000 µm.

**Figure S9:**
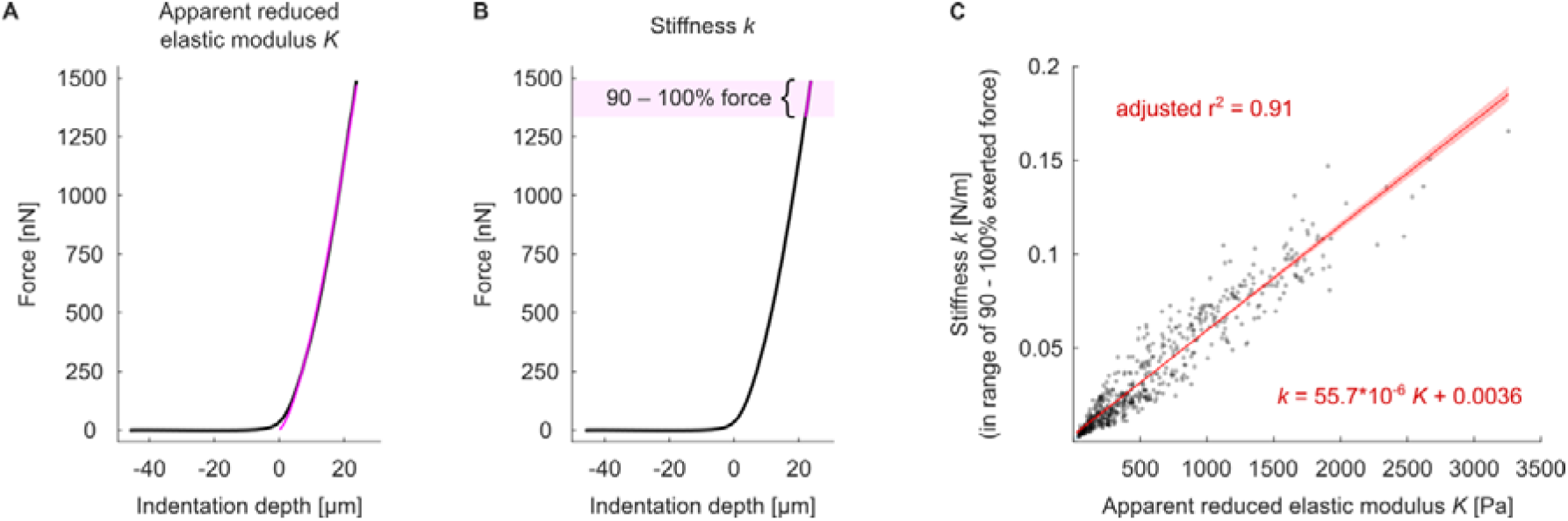
Apparent reduced elastic modulus *K* and stiffness *k* are tightly correlated. Data relate to Figures 1 and 2. AFM force-distance curves were fitted with (**A**) the Sneddon model (pink line) to obtain the apparent reduced elastic modulus *K*, and (**B**) a linear function (pink line) fitted to the range of 90 – 100% of the exerted force (area shaded in light pink) to obtain the stiffness *k* (i.e. the slope of the function). (**A**) and (**B**) show the same example curve. (**C**) Correlation between the apparent reduced elastic moduli *K* and stiffnesses *k*. We found a high degree of positive correlation (Pearson’s correlation coefficient: 0.95), suggesting that both approaches obtain qualitatively similar results. Grey dots represent individual force-distance curves (n = 559 force-distance curves, representing all force-speed combinations from all three anatomical planes; if >10 curves per combination existed, 10 curves were sampled randomly). Red line with shaded red area: Linear regression with 95% confidence interval. Regression parameters and adjusted r^2^ value are indicated in the figure panel.

**Table S1: Number of animals (N) and AFM measurements (n) contributing to grey and white matter measurements at different force-speed combinations in all three anatomical planes.** For every force-speed combination investigated in each anatomical plane in both grey and white matter shown in Figure 2, this table provides the numbers of animals (N), the total number of measurement points (n_total_), the minimum and maximum number of measurement points per animal (n_min_/N and n_max_/N), the animal IDs, the median apparent elastic modulus per animal, and the mean and standard deviation of the median apparent elastic moduli across all investigated animals. The latter two measures have been used for data representation in Figure 2 and Figures S3 and S4. In total, 27,940 AFM measurements contributed to this dataset. Raw data are available online (DOI: <u>10.5281/zenodo.14630529</u>).

**Table S2: Number of animals (N) contributing to the calculation of the grey-to-white-matter elasticity ratio *K_g_/K_w_* at different force-speed combinations in all three anatomical planes.** For every force-speed combination investigated in each anatomical plane shown in Figure 2, this table provides the numbers of animals (N), the animal IDs, the *K_g_/K_w_* ratio for each animal (calculated by dividing the median grey matter apparent elastic modulus by the median white matter apparent elastic modulus), and the mean and standard deviation of the *K_g_/K_w_* ratios across all investigated animals. The latter two measures have been used for data representation in Figure 2 and Figures S3 and S4. Raw data are available online (DOI: <u>10.5281/zenodo.14630529</u>).

